# Spatiotemporal mosaic patterning of pluripotent stem cells using CRISPR interference

**DOI:** 10.1101/252189

**Authors:** Ashley R.G. Libby, David A. Joy, Po-Lin So, Mohammad A. Mandegar, Jonathon M. Muncie, Valerie M. Weaver, Bruce R. Conklin, Todd C. McDevitt

## Abstract

Morphogenesis results from the interactions of asymmetric cell populations to form complex multicellular patterns and structures comprised of distinct cell types. However, current methods to model morphogenic events offer little control over parallel cell type co-emergence and do not offer the capability to selectively perturb gene expression in specific subpopulations of cells. We have developed an *in vitro* system that can spatiotemporally interrogate cell-cell interactions and multicellular organization within human induced pluripotent stem cell (hiPSC) colonies. We examined the effects of independently knocking down molecular regulators of cortical tension and cell-cell adhesion using inducible CRISPRi: Rho-associated kinase-1 (ROCK1) and E-cadherin (CDH1), respectively. Induced mosaic knockdown of ROCK1 or CDH1 in hiPSC populations resulted in differential patterning events within hiPSC colonies indicative of cell-driven population organization. Patterned colonies retained an epithelial phenotype and nuclear expression of pluripotency markers. Gene expression within each of the mixed populations displayed a transient wave of differential expression with induction of knockdown that stabilized in coordination with intrinsic pattern formation. Mosaic patterning of hiPSCs enables the genetic interrogation of emergent multicellular properties of pluripotent cells, leading to a greater mechanistic understanding of the specific molecular pathways regulating the dynamics of symmetry breaking events that transpire during developmental morphogenesis.

**SIGNIFICANCE:** Human embryonic development entails a series of multicellular morphogenic events that lead to primitive tissue formation. Attempts to study human morphogenic processes experimentally have been limited due to divergence from model organisms and the inability of current human *in vitro* models to accurately control the coincident emergence of heterogeneous cell populations in the spatially controlled manner necessary for proper tissue structure. We developed a human induced pluripotent stem cell (iPSC) *in vitro* model that enables temporal control over the emergence of heterotypic subpopulations of cells. We examined mosaic knockdown of two target molecules to create predictable and robust cell-patterning events within hiPSC colonies. This method allows for dynamic interrogation of intrinsic cell mechanisms that initiate symmetry breaking events and provides direct insight(s) into tissue developmental principles.

## BACKGROUND

Early morphogenic tissue development requires the robust coordination of biochemical and biophysical signaling cues that dictate cell-cell communication, multicellular organization, and cell fate determination. (C.A. Burdsal et al., 1993; Leckband et al., 2011; Montero and Heisenberg, 2004). A hallmark of morphogenesis is the asymmetric co-emergence of distinct cell populations that self-organize to form developmental patterns, multicellular structures, and ultimately functional tissues and organs (Lancaster and Knoblich, 2014; Sasai, 2013). For example, during gastrulation, the blastocyst transitions from a relatively homogeneous population of pluripotent cells to a spatially-organized, multicellular composition of distinct progenitor cells. Therefore, in order to study morphogenesis, it is essential to promote the coincident development of analogous heterogeneous populations *ex vivo*. Human pluripotent stem cells (hPSCs) provide an unlimited source of cells that can mimic developmental differentiation processes and maintain the ability to self-organize into tissue-like structures, such as optic cups, gut organoids, or stratified cortical tissues (Eiraku et al., 2008, 2011; Spence et al., 2010). However, due to the intrinsic variability of organoids (Bredenoord et al., 2017) and the lack of alternative human models that faithfully promote asymmetric emergence, many of the mechanisms that control and coordinate morphogenesis remain undefined.

Controlling cellular heterogeneity *in vitro* is often achieved by independent differentiation of hPSCs followed by re-combination of distinct cell types, which fails to mimic parallel cell type emergence (Matthys et al., 2016). Attempts to engineer *in vitro* systems that yield controlled emergence of spatial organization often rely on extrinsic physical restriction of cells to direct subsequent multicellular pattern formation (Hsiao et al., 2009; Warmflash et al., 2014). Physical constraints allow for the observational study of cell-cell interactions within defined regions, but artificially restrict cell behaviors, which can limit the degrees of freedom that can be exerted by cells. Additionally, current tools to elucidate gene function, such as genetic knockouts or siRNA (Boettcher and McManus, 2015), can not selectively perturb gene expression of subpopulations of cells *in situ*, which is required to generate controlled asymmetry analogous to embryonic morphogenesis. Several of these limitations can be addressed with the use of inducible CRISPR interference (CRISPRi) systems in mammalian cells (Larson et al., 2013; Mandegar et al., 2016). Introduction of CRISPRi enables temporal regulation over knockdowns of specific genetic targets with limited off-target effects. Temporal knockdown constraints enable the development of precisely-controlled engineered biological systems that can induce well-defined genetic perturbation at explicit times and within defined populations of cells to mimic developmental symmetry breaking events.

Morphogenic asymmetries arise from reorganization of cells due to local changes in mechanical tissue stiffness and cell adhesions that facilitate physical organization of developing embryos (Krieg et al., 2008; Maître et al., 2012). Mechanical rearrangement is necessary for many aspects of morphogenesis, including cell polarity, collective movement, multicellular organization, and organ size regulation (Arboleda-Estudillo et al., 2010; Maître, 2017). Differential adhesion (Foty and Steinberg, 2004, 2005) and cortical tension (Essen, 1997; Krieg et al., 2008) are critical determinants of mechanically driven cell sorting, where both processes are known to contribute to tissue organization (Lecuit and Lenne, 2007). In cortical tension-dominated sorting, varying actin cytoskeleton-generated cortex tension stimulates sorting of individual cells, while in differential adhesion-directed sorting, the homophilic adhesions between cells promote subpopulation segregation.

In this study we employed an inducible CRISPRi system in induced pluripotent stem cells (hiPSCs) to silence key proteins that regulate cell adhesion and cortical tension. In order to genetically induce symmetry breaking events within human pluripotent stem cell populations, colonies were examined throughout the induction of mosaic KD, enabling the study of how the creation of physical asymmetries in an otherwise relatively homogeneous population leads to multicellular organization and pattern formation. We show that induction of mosaic KD of ROCK1 or CDH1 results in a “bottom-up” cell-driven pattern formation of human PSC colonies while preserving pluripotency.

## RESULTS

### CRISPRi KD in human iPSCs modulates epithelial morphology

ROCK1 and CDH1 were selected as orthogonal gene targets to manipulate human iPSC population organization by changing the intrinsic mechanics of distinct cell populations. ROCK1 regulates actin-myosin dynamics (Fig.1A), which contribute to a cell’s cortical tension (Salbreux et al., 2012). In addition, ROCK inhibition is often used in human pluripotent cell culture and has been implicated in pluripotency maintenance (McBeath et al., 2004; Ohgushi et al., 2015). Similarly, CDH1, a classic type I cadherin adhesion molecule, is widely associated with pluripotency and early morphogenesis (Heasman et al., 1994; Przybyla et al., 2016; Ringwald et al., 1987), and its down-regulation parallels the induction of patterning events via differential adhesion (Fig.1A). To establish an inducible CRISPRi knockdown (KD) of ROCK1 or CDH1, we used a doxycycline (DOX)-inducible CRISPRi iPSC line (CRISPRi-Gen1C) (Mandegar et al., 2016)(Fig. 1B). Guide RNA (gRNA) sequences designed to target the transcription start site of ROCK1 or CDH1 (Sup. Table 1) were introduced into CRISPR-Gen1C hiPSCs and KD was induced by the addition of DOX (2μM) into cell culture media. 3 days of KD induction resulted in <30% of ROCK1 mRNA levels compared to hiPSCs without DOX treatment and <10% CDH1 mRNA in CDH1 KD hiPSCs compared to untreated controls (Fig. 1C). Protein KD followed a similar trend where KD populations compared to untreated controls resulted in <20% ROCK1 protein and <10% of CDH1 protein by day 3 of DOX treatment, and reduced protein levels were maintained through day 6 of CRISPRi induction (Fig.1C, Sup. Fig.1).

**Figure 1:**
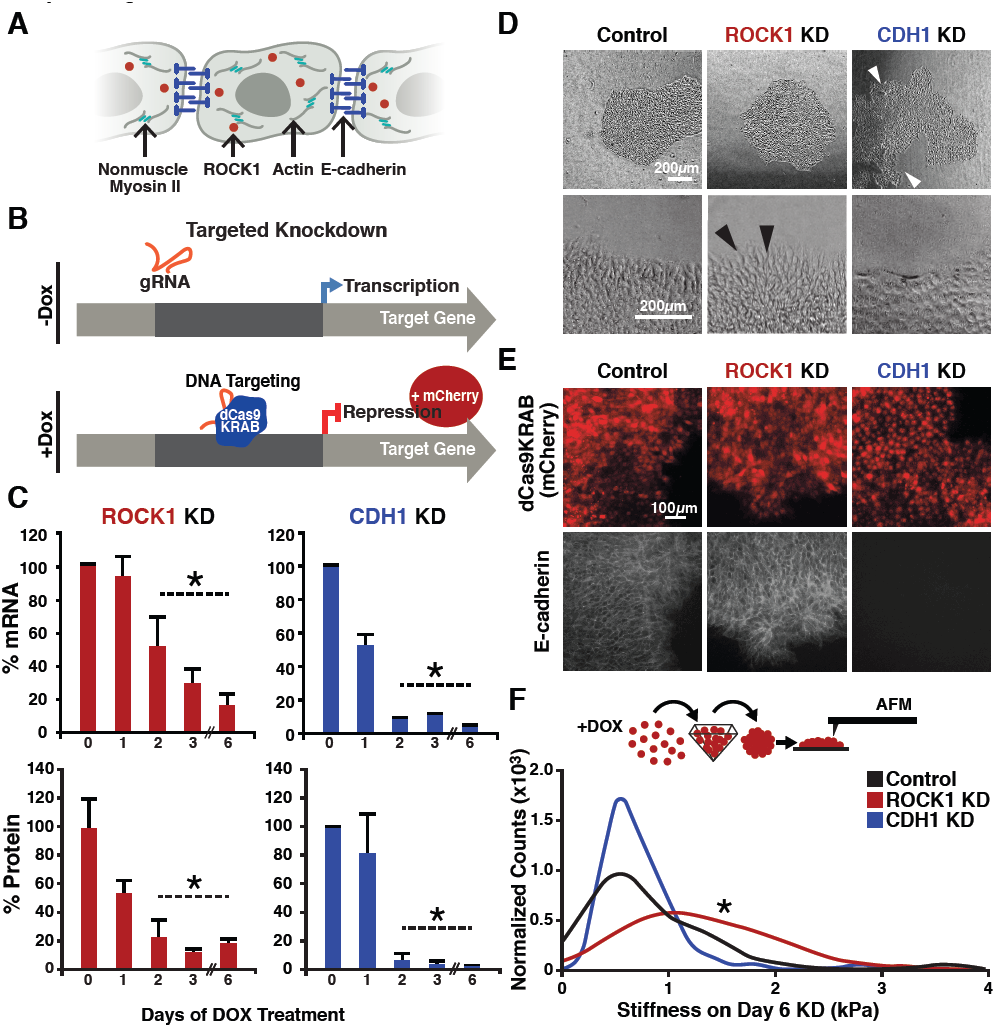
CRISPRi of ROCK1 and CDH1 modulate physical properties of the cell. (A) Schematic of ROCK1 and CDH1 within a cell. CDH1 is a trans-membrane adhesion molecule that locates to the borders of cells and ROCK1 is a cytoplasmic kinase that acts upon non-muscle myosin II. (B) Schematic of the CRISPRi system. Doxycycline addition to the hiPSC culture media leads to the expression of mCherry and dCas9-KRAB to induce knockdown of target gene. (C) qPCR and western blot quantification of knockdown timing; knockdown of both mRNA and protein were achieved by day 3 of DOX treatment when compared to untreated hiPSCs (P < 0.05, n=3, data represent mean ± SD). (D) Brightfield imaging of knockdown hiPSCs indicated morphological differences in colony shape (white arrows) and cell extensions (black arrows) at colony borders. (E) Live-reporter fluorescence for dCas9-KRAB expression (red) and immunostaining for CDH1 (gray) demonstrated loss of CDH1 in induced ECAD CRISPRi hiPSCs, but maintenance of CDH1 contacts in the off-target control and ROCK1 KD hiPSCs. (F) Atomic force microscopy (AFM) of knockdown populations exhibited a two-fold increase in Young’s elastic modulus of ROCK knockdown cells compared to control and CDH1 knockdown cells (P < 0.05, n = 36, 65, 72 force point for Control, ROCK1 KD, and CDH1 KD, respectively, area under curve = 1).

Both the ROCK1 KD cells and the CDH1 KD cells retained epithelial hiPSC morphologies without single cell migration away from the colonies (Fig. 1D). However, CDH1 KD hiPSCs displayed irregular colony shapes that maintained smooth peripheral edges, but often contained regions lacking cells within colonies (Fig. 1D). Conversely, ROCK1 KD hiPSCs displayed round colony shapes (similar to wildtype hiPSCs) but individual cells along the border of ROCK1 KD colonies extended protrusions out away from the colony (Fig. 1D). hiPSCs treated with the small molecule ROCK inhibitor Y-27632 yielded a similar morphology to the ROCK1 KD hiPSCs with extended cell protrusions at the colony borders (Sup. Fig. 2).

To further confirm the physical effects of knocking down CDH1 or ROCK1 selectively in hiPSCs, we performed immunofluorescent (IF) staining of CDH1 expression. After 5 days of DOX treatment, CDH1 KD hiPSCs exhibited a complete loss of CDH1 expression, as expected, while the ROCK1 KD hiPSCs and the control hiPSCs (with off-target CRISPRi guide) maintained robust expression of CDH1 along the plasma membrane (Fig.1E). To interrogate cell cortical tension, the contact angles between cells were measured based upon IF of zona occluden-1 (ZO1), a protein associated with tight junctions (Sup. Fig. 3A). Contact angles were not statistically different in either the ROCK1 KD or CDH1 KD cells compared to the control, however all populations displayed a subtle DOX effect that was not significantly different between any of the groups (Sup. Fig. 3B). However, when direct measurements of hiPSC elasticity were taken using atomic force microscopy after 6 days of KD, ROCK1 KD cells displayed a two-fold higher cortical stiffness than the control and CDH1 KD populations, whereas the latter groups did not differ from one another (Fig. 1F). Therefore, CRISPRi silencing of targeted genes associated with cellular mechanical properties resulted in distinct physical differences between the otherwise similar cell populations.

### Mosaic CRISPRi silencing results in multicellular organization

To examine whether mosaic KD of a single molecule impacted hiPSC organization, ROCK1 or CDH1 CRISPRi populations were pretreated with DOX for 5 days and mixed with isogenic wildtype hiPSCs that constitutively expressed GFP (WT-GFP) at a 1:3 ratio. Forced aggregation of ROCK1 KD:WT-GFP hiPSCs or CDH1 KD: WT-GFP hiPSCs and subsequent re-plating was used to create individual colonies of randomly mixed ROCK1 KD hiPSCs or CDH1 KD hiPSCs with the WT-GFP cells (Fig. 2A). After 5 days in mixed culture, ROCK1 KD cells sorted radially from the WT-GFP cells, clustering primarily at the edges of the colonies (Fig. 2B). However, separation of the ROCK1 KD cells did not result in distinct smooth borders between the WT-GFP and ROCK1 KD hiPSC populations. In contrast, CDH1 KD cells robustly separated from the GFP-WT population, forming sharp boundaries between populations irrespective of their spatial location within the colony (Fig. 2B).

**Figure 2:**
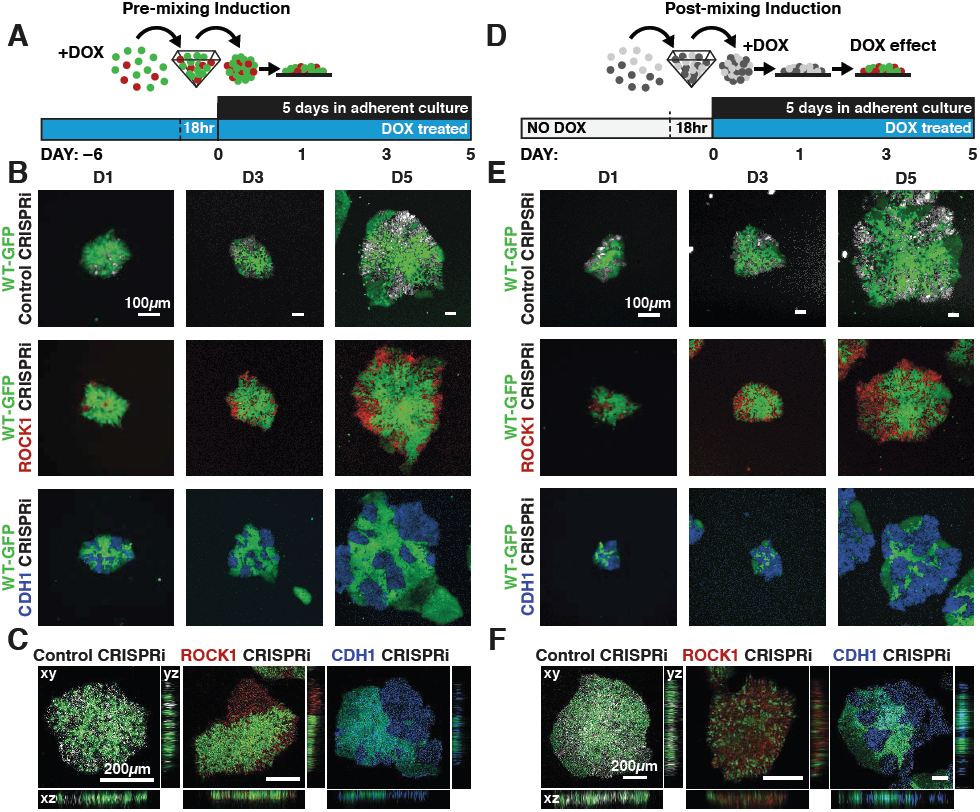
Cell autonomous pattern emergence in mixed population colonies. (A) Schematic of experimental timeline. WT-GFP and ROCK1‐ or ECAD CRISPRi hiPSCs were pretreated with doxycycline for 6 days before aggregation in pyramidal microwells and re-plating as mixed colonies. (B) Live cell imaging of pattern emergence over time from mixing colonies. Control populations remain mixed, ROCK1 KD hiPSCs cluster radially at borders of colonies, and CDH1 KD populations sort themselves from WT-GFP hiPSCs regardless of location within colony (C) Confocal microscopy of patterned colonies of hiPSCs with KD induction prior to mixing. (D) Schematic of experimental timeline for WT-GFP and ROCK1‐ or ECAD CRISPRi hiPSCs treated with doxycycline upon re-plating as mixed colonies. (E) Live cell imaging of pattern emergence in post-mixing induction colonies, where CRISPRi KD is induced after cell population mixing. (F) Confocal microscopy of patterned hiPSC colonies with KD induction upon mixing populations, where ROCK KD cells stack vertically with WT-GFP hiPSCs.

To determine whether pattern emergence was impacted by the relative proportion of mosaic KD within a colony, KD cells were mixed with control CRISPRi hiPSCs lacking any gRNA or fluorescent protein at varying cell ratios of 1:1, 1:3, and 3:1. Clustering of ROCK1 KD cells was less apparent as the proportion of ROCK1 KD cells within a colony increased. In fact, increasing ROCK1 KD hiPSCs to 75% of the colony resulted in the entire colony morphology displaying characteristics of a pure ROCK1 KD colony (Sup. Fig.4A). On the other hand, the CDH1 CRISPRi cells separated from the colorless CRISPRi population, irrespective of cell ratio, indicating that the spatial organization occurred regardless of relative population size within a hiPSC colony. The ability of both the ROCK1 and CDH1 CRISPRi KD populations to physically partition from otherwise identical CRISPRi engineered hiPSCs that lacked a gRNA confirms that the production of dCas9-KRAB is not responsible for the previously observed pattern formation when CRISPRi KD hiPSCs were mixed with the WTGFP cells, but rather that the segregation is a direct result of KD of the target gene (Sup. Fig.4A).

Based on the sorting behaviors of ROCK1 KD:WT-GFP and CDH1 KD:WT-GFP colonies when the KD of ROCK1 or CDH1 was present at the time of mixing, we next examined whether induction of mosaic KD after mixing resulted in similar sorting patterns as previously observed. This scenario more accurately models the onset of initial symmetry breaking events among homogeneous pluripotent cells during embryonic development. Non-induced CRISPRi populations were mixed with WT-GFP hiPSCs (1:3 ratio) and upon replating, treated with DOX to induce KD (Fig. 2D). ROCK1 KD post-mixing within mosaic colonies did not result in noticeable radial segregation of ROCK1 KD hiPSCs from WT-GFP hiPSCs (Fig. 2E), as observed for premixed colonies. Instead, the post-mixing ROCK1 mosaic KD colonies exhibited greater vertical stacking of ROCK1 KD cells and WT-GFP cells in the z-plane of the mixed colonies (Fig. 2F), whereas the pre-induced mixed colonies remained segregated primarily in a 2D planar manner (Fig. 2C). In contrast, the mosaic silencing of CDH1 post-mixing maintained robust segregation of the CDH1 KD cells from the WT-GFP hiPSCs, although the borders between cell populations lacking CDH1 contacts and neighboring WT-GFP cells were somewhat less distinct than the pre-induced CDH1 KD:WT-GFP mixed colonies. Overall, the inducible CRISPRi mixed colonies displayed the ability to mimic several different types of intrinsic symmetry breaking events that resulted in distinct cell sorting and multicellular pattern formation.

### Mosaic human iPSC colonies retain a pluripotent phenotype

Colony morphology and expression of epithelial markers, such as epithelial cell adhesion molecule (EpCAM), were examined to determine if the cells that lost CDH1 expression segregated from their WT-GFP neighbors due to delamination, or loss of the epithelial phenotype characteristic of hiPSCs. ROCK1 KD:WT-GFP and CDH1 KD:WT-GFP colonies maintained an epithelial morphology throughout 6 days of CRISPRi silencing (Fig. 3B) with no observed migration by CRISPRi-modulated cells away from the colonies. Both ROCK1 KD and CDH1 KD hiPSCs within mixed colonies expressed EpCAM at cell-cell boundaries after 6 days of CRISPRi induction despite changes in cortical tension or intercellular adhesion due to loss of ROCK1 or CDH1, respectively (Fig. 3B). Furthermore, ROCK1/CDH1 KD hiPSCs displayed cell junction-localized β-catenin in pure colonies after 6 days of CRISPRi induction, suggesting maintenance of adherens junctions and epithelial colonies (Sup. Fig. 5A).

**Figure 3:**
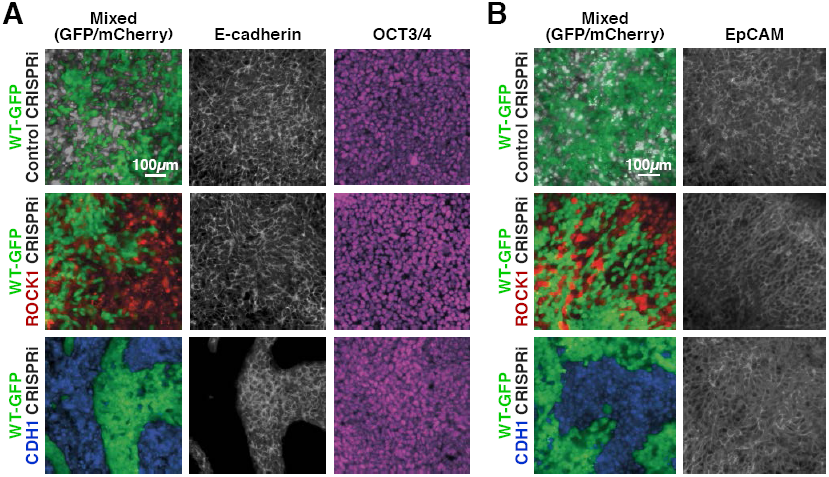
Maintenance of nuclear pluripotency markers and epithelial phenotype. (A) Immunostaining for E-cadherin (CDH1) and OCT3/4 in patterned hiPSC colonies demonstrating nuclear localized OCT3/4 throughout the mixed populations. (B) Immunostaining of EpCAM for mixed colonies displayed relatively uniform expression regardless of KD.

Since the decrease of CDH1 is commonly associated with loss of pluripotency in hPSCs, the expression and localization of the common pluripotency transcription factors, OCT3/4 and SOX2, were examined. Both proteins maintained strong nuclear expression in pure ROCK1 KD or CDH1 KD colonies after 6 days of KD induction (Sup. Fig.6A). Moreover, despite the physical segregation of cells induced by KD in mixed populations, no pattern could be observed based on pluripotency marker expression (Fig. 3A). Furthermore, the abundance of OCT3/4 and SOX2 transcripts was unchanged in pure colonies of CDH1 KD cells and though variable in pure colonies of ROCK1 KD hiPSCs, was not significantly different (Sup. Fig. 6B). These results indicate that the loss of ROCK1 or CDH1 is not sufficient to disrupt the pluripotent gene regulatory network and induce an exit from the pluripotent state.

### Mixed populations of KD cells display transient gene expression changes in coordination with emergence of patterns

Since pluripotency appeared to be maintained irrespective of mosaic patterning, we investigated whether induction of mosaic patterns resulted in gene expression changes over time. We focused on a select set of genes involved in pluripotent stem cell signaling, early lineage fate transitions, and regulation of physical cell properties (Sup. Table 2) in mixed populations at days 1, 3 and 6 after KD induction (Fig. 4A). To take into account potential gene expression changes that result from mixing hiPSC lines, un-induced mixed populations and un-induced pure populations were included as controls. To distinguish any changes that resulted solely from KD of either ROCK1 or CDH1, pure induced populations were also included. Overall there were few changes in gene expression that resulted from mixing un-induced CRISPRi populations with WT-GFP, and therefore subsequent data was normalized to both pure un-induced populations and mixed-uninduced populations to minimize false positives that resulted from mixing of cell lines without induction of knockdown. The ROCK1 KD hiPSCs in the ROCK1 KD:WT-GFP colonies underwent a transient wave of expression changes in genes that regulate pluripotency maintenance. Interestingly, there were some gene expression changes that were specific to mosaic KD within mixed populations. However, by day 6 of ROCK1 KD, gene expression returned to equivalent levels prior to induction (i.e. at day 0) (Fig.4B). Similarly, the CDH1 KD hiPSCs from CDH1 KD:WT-GFP colonies also exhibited a transient wave of gene expression changes upon KD of CDH1 and partitioning of CDH1(+) cells from CDH1(-) cells. After 6 days of KD, the only gene that was significantly differentially expressed in the CDH1 KD population was CDH1 and all other examined genes were unchanged compared to the time matched control populations. This was consistent with our previous observation of the maintenance of pluripotency in the ROCK1 KD:WT-GFP and CDH1 KD:WT-GFP colonies (Fig.3). Furthermore, the recovery of homeostatic gene expression profiles closely followed the dynamics of distinct pattern establishment in the mixed populations.

**Figure 4:**
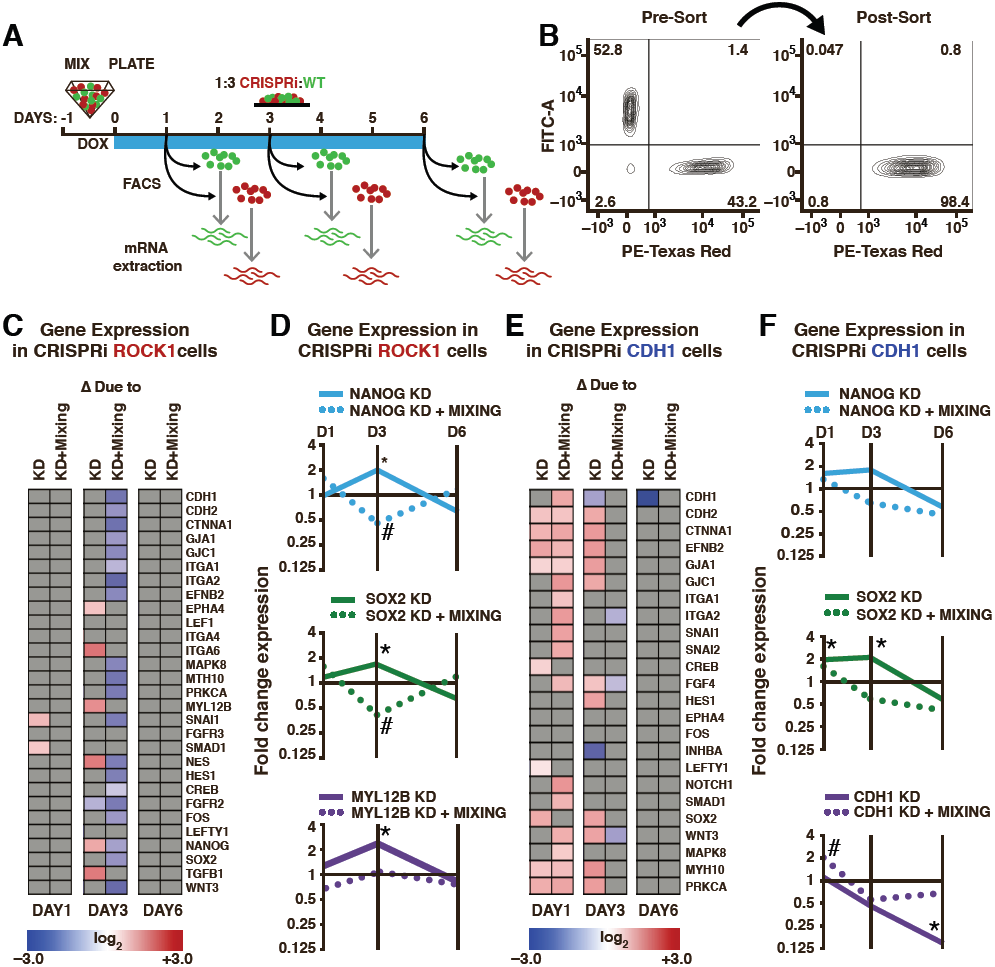
Transient gene expression changes in mixed populations. (A) Schematic of experimental timeline; WT-GFP and ROCK1‐ or ECAD CRISPRi hiPSCs were mixed and re-plated prior to KD induction. Different cell populations were isolated by FACS for mRNA extraction on days 1, 3, and 6 after KD induction. (B) Representative scatter plot of a FACS-sorted population of mCherry+ cells (indicating KD induction) with >98% purity. (C,E) Heat maps display fold-change expression of genes found to display significant changes in ROCK1 or CDH1 KD cells mixed with WT-GFP hiPSCs when compared to time-matched, off-target control hiPSCs. Grey color indicates non-significance. Significance (P < 0.05, n=3) was determined using a one-way analysis of variance (ANOVA) followed by post-hoc pairwise comparisons by Tukey’s tests to determine the affect of mixing populations, the affect of KD, and the affect of KD within a mixed population. (D,F) Plots of specific mRNA expression changes at days 1, 3, and 6. (* and # indicate significance, P < 0.05).

In addition to examining the KD cells, we also examined the gene expression profiles of the neighboring WT cells that constituted the majority of cells in each colony. On day 6 of KD induction, the WT-GFP cells that were mixed with CDH1 KD cells had gene expression patterns that resembled the WT-GFP cells mixed with the control CRISPRi populations, whereas the WT-GFP cells mixed with ROCK1 KD hiPSCs exhibited a different expression profile. Interestingly, the WT-GFP cells mixed with ROCK1 CRISPRi cells demonstrated changes in genes associated with cell sorting and movement, such as ephrins and integrins and up-regulation in myosin proteins (MYH9, MYH10) (Sup. Fig.7B). Overall, the changes in the WT-GFP hiPSC gene expression suggests that targeted manipulation of gene expression in an emerging sub-population can exert non-cell autonomous effects on the opposing population and may be influenced by the respective multicellular organization of the two populations.

## DISCUSSION

In this study we examined the effect of inducing specific genetic KD in subpopulations of hiPSCs within an otherwise homogeneous population of pluripotent cells. Historically, small molecule chemical inhibitors, antibodies, and homogeneous genetic knockouts are often used to interrogate the molecular mechanisms involved in morphogenesis (Lecuit and Lenne, 2007; McBeath et al., 2004; Salbreux et al., 2012). However, these methods can’t selectively discriminate between different cells or they fail to address how the emergence of heterotypic interactions affects cell-cell organization. Here, we report that silencing of target genes by CRISPR interference within only subpopulations provides multiple avenues to genetically control the emergence of asymmetric cell phenotypes and development of multicellular patterns. Specifically, we demonstrate that mosaic KD of target genes, ROCK1 or CDH1, result in distinct patterning events wherein cell driven segregation dictates colony organization without loss of pluripotency (Fig. 5).

**Figure 5:**
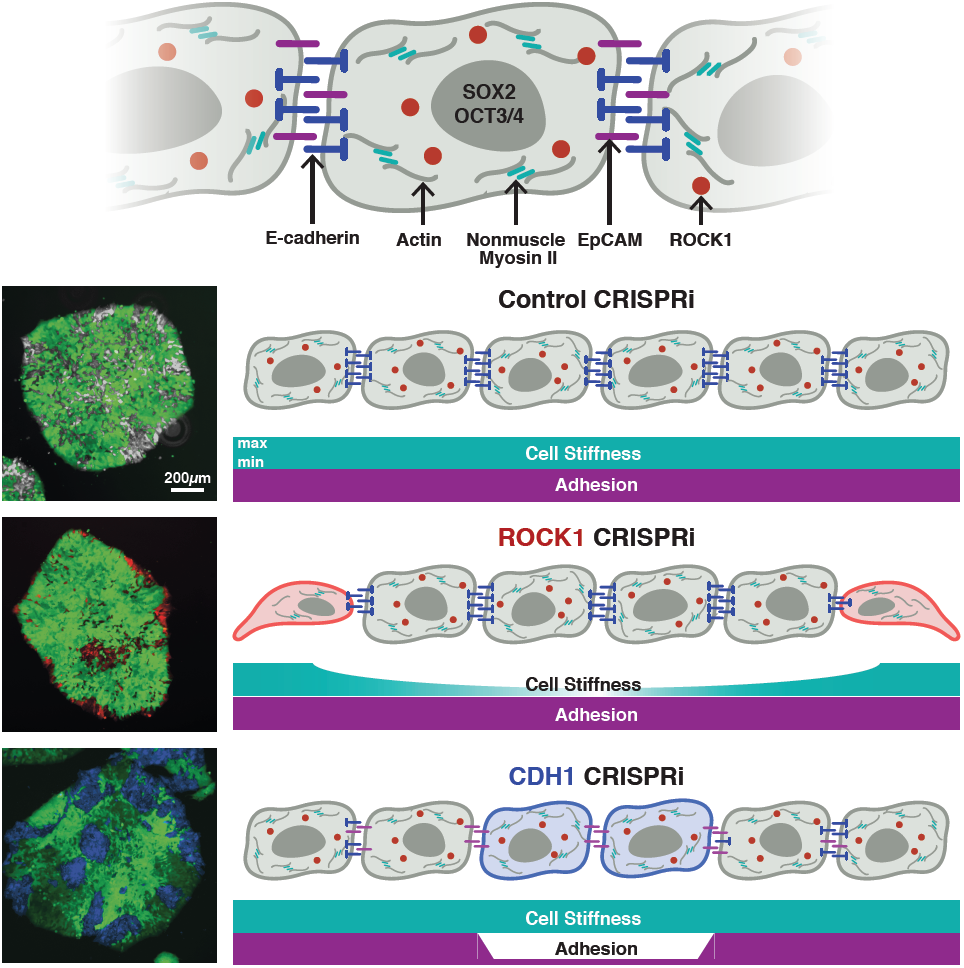
Inducible pattern emergence through the KD of molecules that affect hiPSC physical properties. (A) Schematic of working model of sub-population manipulation where controlled changes in cellular stiffness or cellular adhesion result in specific colony pattern formation. With mosaic KD, ROCK1 produces continuous radial separation of KD cells from WT, where as CDH1 displays discrete islands of KD cells within the WT population.

ROCK1 regulates actin-myosin contraction (McBeath et al., 2004), facilitates expansion of PSCs (Ohgushi et al., 2015; Park et al., 2015), and acute inhibition by small molecules leads to a “relaxed” cell phenotype with decreased stiffness (Kinney et al., 2014; Lee et al., 2006). However, we found that prolonged silencing of ROCK1 in hiPSCs (6 days) resulted in cells that were actually two-fold stiffer than either the CDH1 KD cells or the control CRISPRi cells. The increased cortical stiffness of ROCK1 KD hiPSCs could be due to the difference between the inhibition of an existing protein and KD of the gene. A small molecule inhibitor prevents the function of already existing proteins such that a small amount of functioning protein may escape the inhibitor’s influence. In contrast, CRISPRi only needs to target the ROCK1 gene loci at two alleles to completely abolish protein translation, thus highlighting the strength of genetic perturbation. Additionally, ROCK inhibition is often used as a transient perturbation (24h), whereas long-term KD of ROCK1 (6 days) may induce compensatory effects within the cells that are responsible for the somewhat surprising results. It is likely that long term ROCK1 KD compensation may partially explain why KD of ROCK1 prior to mixing resulted in radially partitioned populations, while post-mixing KD resulted in less segregated populations.

The emergence of autonomous patterning events and separate cell populations is often associated with differentiation, and CDH1 in particular is involved in the control of morphogenesis in a range of species (C.A. Burdsal et al., 1993; Li et al., 2010). Historically, *in vitro* studies of mouse embryonic stem cells often describe CDH1 as a marker of stem cell pluripotency (Li et al., 2012; Soncin and Ward, 2011). However while CDH1 is commonly expressed by pluripotent cells and CDH1 is capable of replacing OCT3/4 during fibroblast reprogramming to pluripotency (Redmer et al., 2011), CDH1 is not essential for maintenance of the pluripotent state (Larue et al., 1996; Soncin et al., 2009; Ying et al., 2008). Our results are consistent with the latter observations where CDH1 KD in hiPSCs did not disrupt the expression of pluripotency markers nor lead to a loss of epithelial phenotype, indicating that KD of CDH1 alone was not sufficient to induce differentiation, but could result in rearrangement of colony structure. Furthermore, the observed maintenance of pluripotency is consistent with recent studies revealing that changes in human CDH1 adhesions coordinate with *in vitro* human stem cell lineage decisions rather than pluripotency maintenance (Przybyla et al., 2016). Changes in CDH1 influencing lineage fate decisions may explain the transient gene expression changes that we observed with the induction of KD in mixed colonies, where the loss of CDH1 potentially primes the cells to respond to a signal for differentiation and without such a signal, the cells return to a ground state of pluripotency. Similar priming has been described in the context of cell-matrix adhesion where differentiation in response to TGFβ signaling is primed by stiffness-dependent integrin signaling (Allen et al., 2012); a similar mechanism may explain the observed transient gene expression changes without loss of pluripotency in CDH1 KD hiPSCs.

The ability to manipulate distinct cell populations allows for robust modeling of human morphogenic events, and thus an expanded understanding of human biology that can be exploited to develop physiologically realistic *in vitro* human tissue models. Cellular location within pluripotent colonies can be thought to parallel the effects seen in early developing blastocysts. Within the early embryo a cell’s location relays signals that dictate initial symmetry breaking events, such as the decision to become trophectoderm instead of inner cell mass. Cells located within the center of an embryo maintain different adhesion contacts (Stephenson et al., 2010) and are subjected to higher tension generated by neighboring cells (Samarage et al., 2015) which then feed back into lineage fate decisions. For example, the Hippo pathway is controlled by a cell’s position within the early blastocyst, where the outer cell layer has the ability to polarize and sequester the signaling molecule angiomotin away from adherens junctions preventing its phosphorylation and activation that would occur in an internal cell that maintained cell-cell contacts on all sides (Hirate et al., 2013). Additionally *in vitro* micro-patterned PSC colonies have been reported to display spatially dependent germ layer patterning upon differentiation (Etoc et al., 2016; Tewary et al., 2017; Warmflash et al., 2014). Therefore, the system described here enables the potential to enhance and expand on these previous complementary studies by allowing for the manipulation of local multicellular neighborhoods through subpopulation organization.

Overall, this study capitalized on the ability of CRISPRi to temporally perturb specific molecular regulators of physical cell properties, such as adhesion and tension that resulted in differing multicellular patterns. Moreover, CRISPRi additionally offers the flexibility to target any gene of interest and timing of KD (Gordon et al., 2016; Mandegar et al., 2016), allowing for the creation of dynamic patterns through transient genetic KD which could be used to pre-pattern PSC colonies in various types of multicellular geometries prior to differentiation. Additionally, the ability to induce molecular asymmetry can be applied to co-differentiation, where the temporal induction of specific heterotypic interactions, such as the presentation of a ligand or receptor, can give rise to the coordinated emergence of 2 (or more) cell types under the same culture conditions. In addition, mosaic induction of KD can be used to examine how signals propagate between cells. For example, interrogating how the networks between cells created by either mechanical (adhesions) or chemical gradients (gap junctions), affect lineage fate decisions. Furthermore, the predictable patterning events and potential for control over co-emergence that we establish in this study could aid the eventual control over morphogenic events in organoid systems. Organoids require coordinated heterotypic interactions in a 3D environment in order to self-organize (Bredenoord et al., 2017; Sasai, 2013); the ability to precisely predict and control the organization of multiple cell types in parallel would significantly improve the reproducibility and robustness of *in vitro* tissue modeling. Ultimately, this study identifies a novel strategy to direct the emergence of heterotypic cell populations to control multicellular organization in pluripotent stem cells, and subsequently facilitates the creation of robust models of morphogenesis necessary for the mechanistic study of human developmental tissue patterning and formation.

## METHODS

### Human iPSC line generation and culture

All work with human iPSC lines was approved by the University of California – San Francisco Human Gamete, Embryo and Stem Cell Research (GESCR) Committee. hiPSCs lines were cultured in feeder-free media conditions on growth factor-reduced Matrigel (BD Biosciences) and fed daily with mTeSR^TM^-1 medium (STEMCELL Technologies)(Ludwig et al., 2006). Accutase (STEMCELL Technologies) was used to dissociate hiPSCs to single cells during passaging. Cells were passaged at a seeding density of 12,000 cells per cm^2^ and the small molecule Rho-associated coiled-coil kinase (ROCK) inhibitor, Y-276932 (10μM; Selleckchem) was added to the media upon passaging to promote survival (Park et al., 2015; Watanabe et al., 2007).

For the generation of the CDH1 CRISPRi line, five CRISPRi gRNAs were designed to bind within 150bp of the TSS of CDH1 and cloned into the gRNA-CKB vector using BsmBI ligation following the previously described protocol (Mandegar et al., 2016)(Sup Table 1). gRNA expression vectors were nucleofected into the CRISPRi-Gen1C hiPSC line from the Conklin Lab using the Human Stem Cell Nucleofector Kit 1 solution with the Amaxa nucleofector 2b device (Lonza). Nucleofected cells were seeded into 3 wells of a 6-well plate (∼7400 cell/cm^2^) in mTeSR^TM^-1 media with Y-27632 (10μM) for two days, and treated with Blasticidin (10μg/ml) for a selection period of 7 days. Surviving colonies were pooled and passaged in mTeSR^TM^-1 with blasticidin and Y-27632 for a single day then transitioned to mTeSR^TM^-1 media only. Once stable polyclonal populations of CDH1 CRISPRi hiPSCs for each of the five guides were established, the cells were incubated with doxycycline (2μM) for 96hrs. KD efficiency was evaluated by mRNA collection and subsequent qPCR, comparing levels of transcript with a time matched control of the same line without CRISPRi induction. The cell line with the guide producing the best KD was selected (gRNA −6).

To generate the WT-GFP line, 2 million WTC clone11 human iPSCs were nucleofected as previously described with the knock-in plasmid containing a CAG promoter-driven EGFP and AAVS1 TALEN pair vectors (Sup. Fig. 8A). After cell recovery, puromycin (0.5μg/ml) was added to the media for a selection period of 7 days. Individual stable EGFP expressing colonies were picked using an EVOS FL microscope (Life Technologies) and transferred to individual wells of a 24 well plate in mTeSR media with Y-27632 (10μM) and subsequently expanded into larger vessels.

All cell lines were karyotyped by Cell Line Genetics and were deemed karyotypically normal before proceeding with experiments (Sup. Fig. 8B)

### Generation of mixed colonies

Cell aggregates of ∼100 cells were created using 400 X 400μm PDMS microwell inserts in 24-well plates (∼975 microwell per well) similar to previously published protocols (Hookway et al., 2016; Ungrin et al., 2008). Dissociated hiPSC cultures were resuspended in mTeSR^TM^-1 supplemented with Y-27632(10μM), mixed at proper ratios and concentration (100 cells/well), added to microwells, and centrifuged (200rcf). After 18 h of formation, 100 cell aggregates were transferred in mTeSR^TM^-1 to Matrigel coated 96 well plates (∼15 aggregates/cm^2^) and allowed to spread into 2D colonies.

### Western blot

Human iPSCs were washed with cold PBS, incubated for 10 min on ice in RIPA Buffer (Sigma-Aldrich), and supernatant collected. The supernatant protein content was determined using a Pierce BCA Protein Assay kit (Thermofisher Scientific) colorimetric reaction and quantified on a SpectraMax i3 Multi-Mode Platform (Molecular Devices). Subsequently, 20μg of protein from each sample was resolved by SDS-PAGE, and transferred to a nitrocellulose membrane (Invitrogen). The membranes were incubated overnight at 4°C with primary antibodies: anti-ROCK1 (AbCAM 1:200), anti-CDH1 (AbCAM 1:200), anti-GAPDH, (Invitrogen 1:10,000), followed by incubation (30 minutes at room temperature) with infrared secondary antibodies: IRDye 800CW and IRDye 680CW (LI-COR 1:13,000), and imaged on the Odyssey Fc Imaging System (LI-COR Biosciences). Protein levels were quantified using Image Studio Lite (LI-COR Biosciences).

### RNA isolation and qPCR

Total RNA isolation was performed using an RNeasy Mini Kit (QIAGEN) according to manufacturer’s instructions and quantified with a Nanodrop 2000c Spectrometer (ThermoFisher). cDNA was synthesized by using an iScript cDNA Synthesis kit (BIORAD) and the reaction was run on a SimpliAmp thermal cycler (Life Technologies). To quantify individual genes, qPCR reactions were run on a StepOnePlus Real-Time PCR system (Applied Biosciences) and detected using Fast SYBR Green Master Mix (ThermoFisher Scientific). Relative gene expression was determined by normalizing to the housekeeping gene 18S rRNA, using the comparative threshold (C_T_) method. Gene expression was displayed as fold change of each sample (ROCK1 CRISPRi or CDH1 CRISPRi) versus the off target guide control (KCNH2 CRISPRi). The primers were designed using the NCBI Primer-BLAST website and are listed in Sup. Table 2. Statistical analysis was conducted using a two-tailed unpaired *t*-test between any two groups (p<0.05, n=3).

### Atomic Force Microscopy

All AFM indentations were performed using an MFP3D-BIO inverted optical atomic force microscope (Asylum Research) mounted on a Nikon TE2000-U inverted fluorescent microscope. Silicon nitride cantilevers were used with spring constants ranging from 0.04 to 0.06 N/m and borosilicate glass spherical tips 5 μm in diameter (Novascan Tech). Each cantilever was calibrated using the thermal oscillation method prior to each experiment. Samples were indented at 1 μm/s loading rate, with a maximum force of 4 nN. Force maps were typically obtained as a 6x6 raster series of indentations utilizing the FMAP function of the IGOR PRO build supplied by Asylum Research, for a total of 36 data points per area of interest measured every 5 microns. Two 5 micron by 5 micron areas of interest were sampled for each sample. The Hertz model was used to determine the elastic modulus of the sample at each point probed. Samples were assumed to be incompressible and a Poisson’s ratio of 0.5 was used in the calculation of the Young’s elastic modulus.

### Time-lapse imaging

Human iPSC colonies were imaged in 96 well plates (ibidi) on an inverted AxioObserver Z1 (Ziess) with an ORCA-Flash4.0 digital CMOS camera (Hamamatsu). Using ZenPro software, colony locations were mapped and a single colony was imaged daily for 6 days. To obtain time-lapse movies, a single colony was imaged over the course of 12 h at a rate of one picture taken every 30 minutes.

### Immunofluorescence staining

Human iPSC colonies were fixed for 30min in 4% paraformaldehyde (VWR) and washed 3X with PBS. Fixed colonies were permeabilized with 0.3% Trition X-100 (Sigma Aldrich) throughout blocking and antibody incubation steps. Samples were incubated in primary antibodies over night at 4°C, subsequently washed with PBS and incubated in secondary antibodies for an hour at room temperature. Primary antibodies used were: anti-OCT4 (SantaCruz 1:400), anti-SOX2 (AbCAM 1:400), anti-Zo1 (LifeTechnologies 1:400), NANOG (AbCAM 1:300), anti-β-catenin (BD Biosciences 1:200), anti-EpCAM (Millipore 1:200). All secondary antibodies were used at 1:1000 and purchased from Life Technologies.

## FACS

Mixed human iPSC populations and pure population controls were dissociated from tissue culture plates and washed 3X with PBS. A LIVE/DEAD stain (ThermoFisher Scientific) was used per manufacture instructions. Prior to sorting, cells were suspended in PBS supplemented with Y-27632(10μM) and kept on ice. A BD FACSAria II cell sorter (BD Biosciences) was used to isolate pure populations of WT-GFP and CRISPRi hiPSCs by first identifying the live cells via the LIVE/DEAD(350) stain and subsequently sorting the mCherry(+) GFP(-) populations from the mCherry(-)GFP(+) populations directly into TRIzol LS Reagent (ThermoFisher Scientifc). Samples were then stored at −80°C until subsequent mRNA extraction.

### Fluidigm 96.96 Array

Sorted hiPSCs stored in TRIzol LS were thawed on ice and mRNA was extracted using a Direct-zol RNA MiniPrep Plus kit (ZYMO Research) following the manufacturer’s instructions. RNA was converted to cDNA using the iScript cDNA synthesis kit (Bio-Rad). Forward and reverse primers for genes were designed using NCBI’s Primer-BLAST (Sup. Table 3). Primers were pooled to 500nM to enable specific-target amplification and cDNA was amplified with PreAmp Master Mix (Fluidigm) and pooled primers for 15 cycles. Pre-amplified samples mixed with 2X SsoFast EvaGreen Supermix with low ROX(Bio-Rad) and 20X DNA Binding Dye Sample Loading Reagent (Fluidigm), and 10μM primer sets were mixed with 2X Assay Loading Reagent (Fluidigm). 5μl of diluted cDNA and primers and were loaded onto the IFC chip per manufacturer’s instructions and loaded into the chip using the IFC Controller HX (Fluidigm). qPCR was run for 40 cycles in the IFC chip using the BioMark HD in the BioMark HD Data Collection Software. Resulting data was analyzed in the Real-Time PCR Analysis Software. All instruments and software involved with the IFC chip were manufactured by Fluidigm. Gene expression levels were calculated with respect to time matched pure populations of WT hiPSCs, and hierarchically clustered and plotted using Genesis software (Institute for Genomics and Bioinformatics, Graz University of Technology).

### Statistics

Unpaired T-tests were used to compare two groups. One-way analysis of variance (ANOVA) was used to compare three or more groups, followed by post-hoc pairwise comparisons by Tukey’s tests. In gene expression analysis heat maps, all gene expression was normalized to control mixed populations (off target guide without knockdown) to control for any gene expression changes due to mixing or the process of FACS sorting. In all comparisons, significance was specified as p ≤ 0.05.

## ACKNOWLEDGEMENTS

We thank the Gladstone Light Microscopy and Histology Core, the Gladstone Flow Cytometry Core (NIH P30 AI027763, NIH S10 RR028962, and the James B. Pendleton Charitable Trust), and the Gladstone Stem Cell Core for their support and experimental expertise. B.R.C and T.C.M were supported by the Gladstone Institutes. T.C.M. received support from the California Institute of Regenerative Medicine (LA1_C14-08015) and the National Science Foundation (CBET 0939511). B.R.C received support from the National Institutes of Health (U01HL100406, P01HL089707, R01HL130533). We are grateful to Ariel Kauss for providing critical analysis and feedback.

**CONTRIBUTIONS**
A.R.G.L, B.R.C., and T.C.M. conceived the project. A.R.G.L., P.L.S., and T.C.M. designed the experiments. The majority of experiments were performed by A.R.G.L with help from D.A.J. M.A.M. generated the WT-GFP hiPSC line. J.M.M. and V.M.W. provided atomic force microscopy techniques and expertise. A.R.G.L., D.A.J., J.M.M. and T.C.M. analyzed data. A.R.G.L., P.L.S., and T.C.M. wrote the paper with editing from all other authors.

## Notes

**CONFLICT OF INTEREST:** B.R.C. is on the Scientific Advisory Board of CODA Therapeutics making therapies related to pain relief, Cellogy Inc. a cell dynamics measurement company, as well as Scientist.com a scientific services company. B.R.C. is a founder of Tenaya Therapeutics (https://www.tenayatherapeutics.com), a company focused on finding treatments for heart failure, including the use of CRISPR interference to interrogate genetic cardiomyopathies. B.R.C. holds equity in Tenaya, and Tenaya provides research support for heart failure related research. M.A.M. is currently an employee of Tenaya Therapeutics. T. C. M. is a consultant for Tenaya Therapeutics.

